# Lineage and stage-specific expressed *CYCD7;1* coordinates the single symmetric division that creates stomatal guard cells

**DOI:** 10.1101/207837

**Authors:** Annika K. Weimer, Juliana L. Matos, Walter Dewitte, James A.H. Murray, Dominique C. Bergmann

**Affiliations:** Department of Biology, Stanford University, Stanford, CA, USA; Cardiff School of Bioscience, Cardiff University, Wales, United Kingdom; Institute of Biotechnology, University of Cambridge, United Kingdom; Howard Hughes Medical Institute (HHMI), Stanford University, Stanford, CA, USA

**Keywords:** stomatal development, cell cycle, cyclin, differentiation, guard cell

## Abstract

Plants, with cells fixed in place by rigid walls, often utilize spatial and temporally distinct cell division programs to organize and maintain organs. This leads to the question of how developmental regulators interact with the cell cycle machinery to link cell division events with particular developmental trajectories. In *Arabidopsis* leaves, the development of stomata, two-celled epidermal valves that mediate plant-atmosphere gas exchange, relies on a series of oriented stem-cell-like asymmetric divisions followed by a single symmetric division. The stomatal lineage is embedded in a tissue whose cells transition from proliferation to post-mitotic differentiation earlier, necessitating stomatal lineage-specific factors to prolong competence to divide. We show that the D-type cyclin, CYCD7;1 is specifically expressed just prior to the symmetric guard-cell forming division, and that it is limiting for this division. Further, we find that CYCD7;1 is capable of promoting divisions in multiple contexts, likely through RBR-dependent promotion of the G1/S transition, but that CYCD7;1 is regulated at the transcriptional level by cell-type specific transcription factors that confine its expression to the appropriate developmental window.

## Introduction

Development of multicellular organisms requires the coordination and control of cell proliferation with differentiation programs to generate distinct cell types, tissues and organs. Different cell lineages are specified by sets of developmental regulators and display various cell proliferation dynamics, suggesting that the cell cycle machinery might not always be comprised of the same components or controlled in the same way. In *Arabidopsis*, the mature leaf epidermis contains pavement cells, trichomes and stomata, three different functional cell types with their own developmental trajectories. Trichome precursors are specified early and patterned via lateral inhibition networks (Schellmann et al., 2002), and their maturation requires a shift from mitotic to endoreplication programs (Bramsiepe et al., 2010). Pavement cells also endoreplicate as they acquire their lobed morphologies (Katagiri et al., 2016).

Stomata, pivotal for gas exchange between the plant and the environment, are derived from protodermal cells in a process that requires them to first become self-renewing and multi-potent, but then to navigate an ordered set of divisions and differentiation programs to create the mature stoma (Matos and Bergmann, 2014). Stomatal development requires three essential, stage-specific, basic-helix loop-helix (bHLH) transcription factors, SPEECHLESS (SPCH), MUTE and FAMA and their broadly expressed heterodimer partners SCRM/ICE1 and SCRM2 (Kanaoka et al., 2008) (Fig 1A). SPCH drives asymmetric cell divisions that initiate the lineage, creating meristemoids (M) that may undergo continued self-renewing divisions. Plants lacking *SPCH* have no stomatal lineage. MUTE is essential to terminate the asymmetric self-renewing divisions and to induce the differentiation of meristemoids into guard mother cells (GMCs) (MacAlister et al., 2007; Pillitteri et al., 2007); loss of *MUTE* results in excess meristemoids at the expense of GMCs (MacAlister et al., 2007; Pillitteri and Torii, 2007). FAMA is required for the establishment of GCs but also to restrict GMCs to a single division. *fama* mutants exhibit numerous rounds of symmetric and parallel GMC divisions without acquisition of terminal GC identities (Matos et al., 2014; Ohashi-Ito and Bergmann, 2006). Plants bearing mutations in two R2R3 MYB transcription factor genes *FOUR LIPS (FLP)* and *MYB88* also exhibit *fama-like* GMC over-proliferation phenotypes (Lai et al., 2005; Xie et al., 2010).

**Figure 1:**
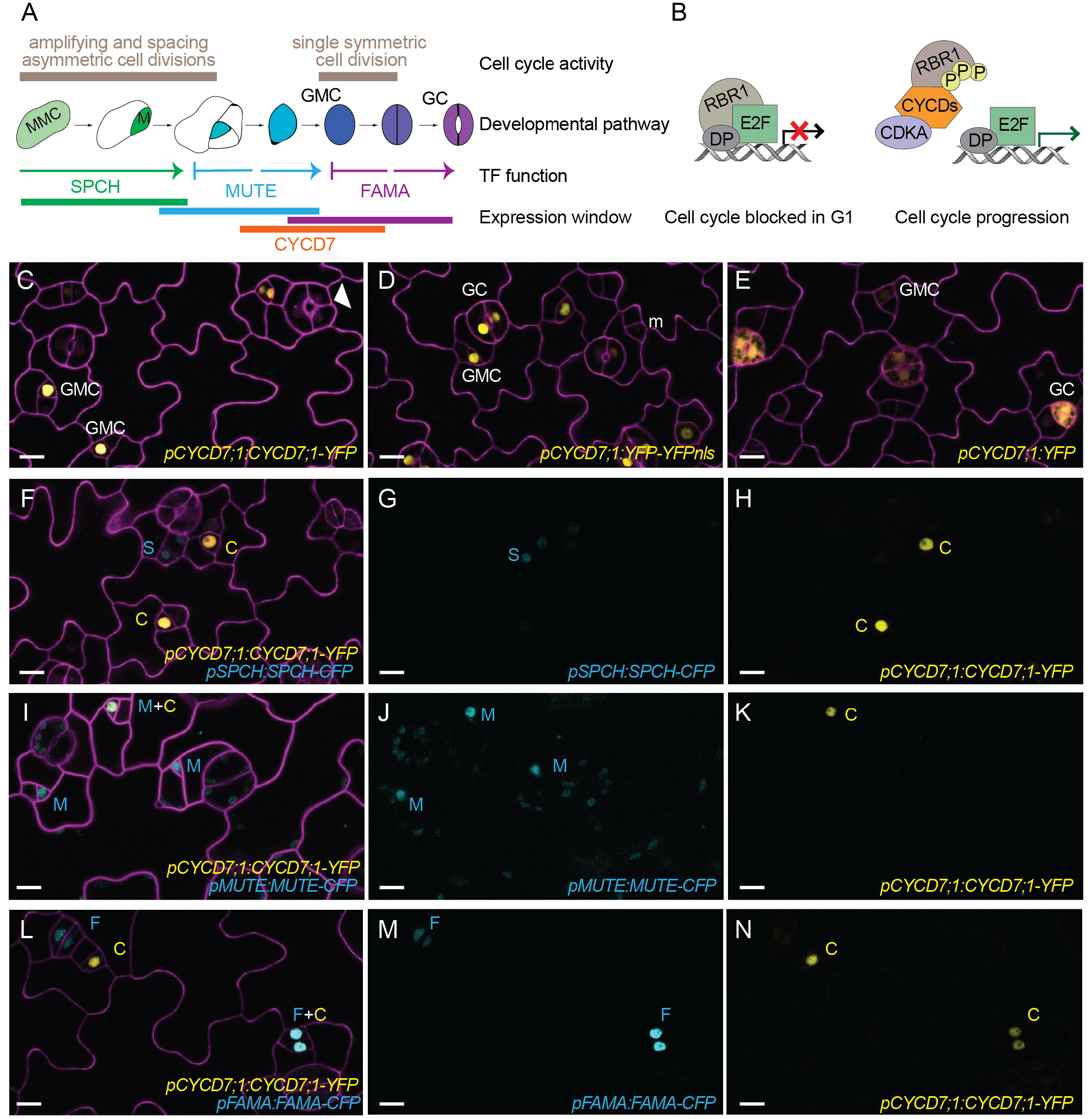
CYCD7;1 is expressed in GMCs prior to the last symmetric division of the stomatal lineage. **(A)** Scheme of stomatal development in *Arabidopsis thaliana*. Cell cycle activity depicted in beige, with cell fate transitions, function and expression window of master bHLH transcription factors SPCH (green), MUTE (blue), and FAMA (purple) and CYCD7;1 (orange). Meristemoid mother cells (lightgreen, MMC) divide asymmetrically to enter the lineage. Meristemoids (green) can undergo amplifying and spacing asymmetric cell divisions until activity is terminated. Guard mother cells (GMC, blue) reenter the cell cycle only once to generate the pair of symmetric guard cells (GC, purple). **(B)** Cartoon of plant RBR1/CYCD complexes driving the G1 to S transition and commitment to divide. RBR1 binds to E2F-DP transcription factors and blocks their ability to induce transcription of S phase genes. CYCDs interact with RBR1 through their L×C×E motif and facilitate phosphorylation of RBR1 by the CDKA;1/CYCD complex. Upon phosphorylation RBR1 releases E2F transcription factors, which leads to expression of S phase genes for DNA replication. **(C-E)** Expression of the translational reporter *pCYCD7;1:CYCD7;1-YFP*, the transcriptional reporters *pCYCD7;1:YFP-YFPnls* and *pCYCD7;1:YFP* (all yellow) in abaxial cotyledons. White arrowheads point at ectopic cell divisions. **(F-N)** Coexpression of *pCYCD7;1:CYCD7;1-YFP* (yellow, C) and *pSPCH:SPCH-CFP* (cyan, S), *pMUTE:MUTE-CFP* (cyan, M) and *pFAMA:FAMA-CFP* (cyan, F). Confocal images were taken at 5 dag (days after germination). Cell outlines (magenta) are visualized with propidium iodide. All images are at the same magnification and scale bar is 10μM.

The varied trajectories of epidermal cells have been useful tools for dissecting cell cycle behaviors. The components of the core cell cycle machinery are highly conserved among eukaryotes, though there has been a large expansion of genes in plants (Harashima et al., 2013; Inzé and De Veylder, 2006). The plant cell cycle is regulated by 5 main cyclin-dependent kinases (CDKs), CDKA;1, CDKB1;1, CDKB1;2, CDKB2;1 and CDKB2;2. CDKs require cyclins (CYC) as binding partners for their kinase activity toward downstream phosphorylation targets. Plants genomes encode much larger families of cyclin genes than animals; for example, *Arabidopsis* encodes at least 32 cyclins (Vandepoele et al., 2002; Wang et al., 2004) and it has been speculated that this expansion allows plants to specifically regulate their postembryonic development (De Veylder et al., 2007; Harashima et al., 2013; Inzé and De Veylder, 2006). D-type cyclins as partners of CDKA;1 are critical for the G1/S cell cycle transition and commitment to divide (Dewitte et al., 2007; Harbour and Dean, 2000; Riou-Khamlichi et al., 2000). Eight out of ten plant CYCDs have an RBR1-binding motif (L×C×E) (Kono et al., 2007; Menges et al., 2003). RBR1, the *Arabidopsis* homolog of the human tumor suppressor protein Retinoblastoma, is crucial for the negative control of the cell cycle at G1/S transition (Desvoyes et al., 2006; Gutzat et al., 2012; Nowack et al., 2012; Uemukai et al., 2005; Zhao et al., 2012). Phosphorylation of RBR1 by CDKA;1/CYCD complexes inactivates its suppression of E2F transcription factors, allowing entry into S phase and commitment to divide (Fig. 1B) (Harashima et al., 2013; Nakagami et al., 2002; Nowack et al., 2012; Umen and Goodenough, 2001).

Here we show how the cell cycle and cell fate transition from GMCs to GCs is regulated by the stomatal-lineage specific G1-S phase cell cycle regulator CYCD7;1. We demonstrate that CYCD7;1 activity is that of a typical D-type cyclin, but its expression window is narrowed by stomatal lineage specific transcription factors. By examining how CYCD7;1 works with the core cell-cycle machinery and with stomatal regulators, and by revealing the phenotypes upon loss and gain of *CYCD7;1* function, we link a core cell-cycle regulator with a specific differentiation process and show how a formative division is initiated but also restricted to allow “one and only one division” in GMCs to create a physiologically functional valve structure from its two identical daughters.

## Results

### CYCD7;1 is expressed prior to the last symmetric division in the stomatal lineage

Among the 10 known D-type cyclins in *Arabidopsis, CYCD7;1* was uniquely enriched in transcriptional profiles of Fluorescence Activated Cell Sorting (FACS) isolated cells of the late stomatal lineage (Adrian et al., 2015). We confirmed this predicted expression in GMCs with transcriptional and translational reporters (Fig. 1C-E) and observed that additional copies of *CYCD7;1-YFP* could force ectopic divisions in GCs, suggesting that the protein could play a role in regulating this division (Fig. 1C, white arrowhead). A translational reporter, *pCYCD7;1:CYCD7;1-YFP,* was characterized previously as peaking in GMCs (Adrian et al., 2015); however, the identity of CYCD7;1 expressing cells was only assessed by morphology. To refine the expression pattern, we co-expressed *pCYCD7;1:CYCD7;1-YFP* with CFP reporters for SPCH, MUTE and FAMA (Fig. 1F-N). SPCH-CFP and CYCD7;1-YFP expression appear to be mutually exclusive, suggesting that CYCD7;1 is not expressed in meristemoids (Fig. 1F-H). MUTE-CFP and CYCD7;1-YFP overlap in some cells, but we also see cells expressing only MUTE or only CYCD7;1. Cells that only express MUTE had the morphology typical of meristemoids, suggesting that MUTE is expressed before CYCD7;1 (Fig. 1I-K). When compared to FAMA expression, CYCD7;1-YFP appears to be expressed before FAMA-CFP in GMCs, briefly together with FAMA in newly divided GCs, and then disappears before FAMA in GCs (Fig. 1L-N). Thus, the expression of CYCD7;1 in the stomatal lineage is temporally and spatially controlled and starts after MUTE expression and finishes before FAMA expression (Fig. 1A).

We did not observe expression of CYCD7;1-YFP in any vegetative tissue from the seedling stage through flowering (data not shown). In adult plants, CYCD7;1-YFP was expressed in pollen sperm cells at anthesis, but not in the vegetative nucleus (Fig. S1). The expression of a D-type cyclin (typically expressed at G1/S) is consistent with the observations that sperm cells undergo an extended S phase in mature pollen grains (Friedman, 1999; Zhao et al., 2012).

Why does CYCD7;1 have such a restricted expression pattern in the stomatal lineage? One possible explanation is that CYCD7;1 has a unique function in GMC divisions. A second possibility is that CYCD7;1 has a canonical role, i.e. it acts like other cyclins in promoting cell divisions, but it is important to be able to tightly control deployment of that role in the stomatal lineage. To distinguish between these models, we characterized plants missing or misexpressing *CYCD7;1,* tested relationships between CYCD7;1 and other cell cycle regulators, and defined how *CYCD7;1* expression was constrained by stomatal lineage transcription factors.

### Ectopic expression of CYCD7;1 triggers divisions while *cycd7;1* mutants decelerate GMC divisions

If CYCD7;1 has canonical CYCD activity, it should be able to promote cell divisions outside its normal expression window. To test this, we expressed CYCD7;1 and CYCD7;1-YFP with the pan-epidermal promoter, ML1 (Roeder et al., 2010). Ectopic expression of CYCD7;1 (YFP-tagged or untagged) induced cell divisions of pavement cells in the leaf (Fig. 2A-C) indicating that CYCD7;1 can function as a canonical D-type cyclin.

**Figure 2:**
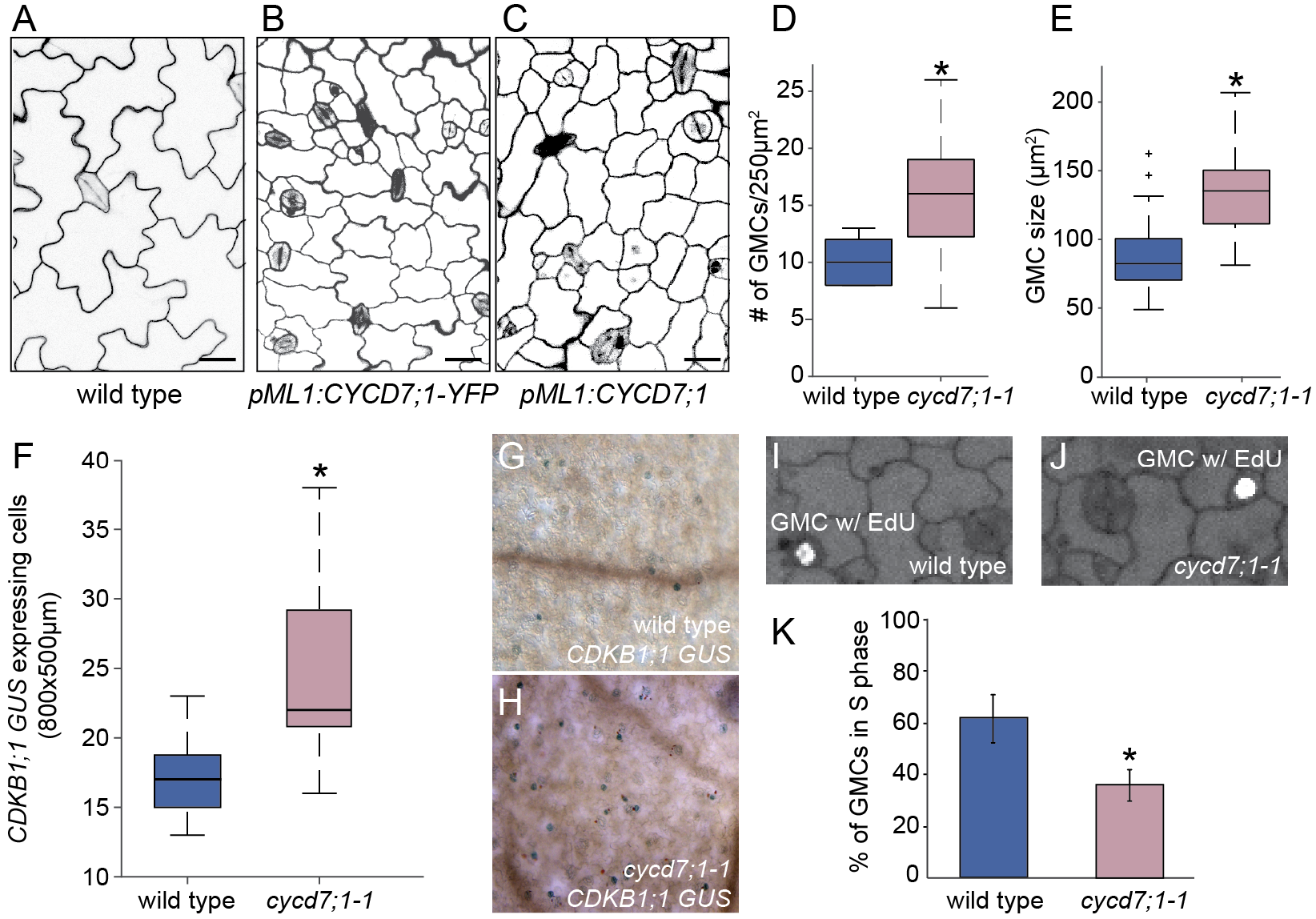
CYCD7;1 promotes cell divisions. **(A-C)** Confocal images of adaxial cotyledon epidermes of wild type, and plant expressing *pML1:CYCD7;1-YFP* and *pML1:CYCD7;1* at 6 dag. Cell outlines were visualized with propidium iodide (magenta). Scale bar 20μM. **(D)** Quantification of the number of GMCs in wild type and *cycd7;1-1* cotyledons at 4 dag. Asterisk indicates significant difference (p-value = 0.0032; Mann-Whitney U test). **(E)** Quantification of GMC area in wild type (N=55) and *cycd7;1-1* (N=51) cotyledons at 4 dag. Asterisk indicates significant difference (p-value = 6.76E-13; Mann-Whitney U test). **(F)** Quantification of cells expressing the *CDKB1;1-GUS* marker in wild type and *cycd7;1-1* cotyledons at 5 dag. Asterisk indicates significant difference (p-value = 0.0023; Mann-Whitney U test). **(G)** Image of wild type cotyledon expressing *CDKB1;1-GUS* marker at 5 dag. **(H)** Image of *cycd7;1-1* cotyledon expressing *CDKB1;1-GUS* marker at 5 dag. **(I)** Image of wild type GMC with EdU (5-ethynyl-2’-deoxyuridine) labeling at 4 dag cotyledon. **(J)** Image of *cycd7;1-1* GMC with EdU labeling, 4-day old cotyledon. (K) Quantification of EdU labeling in wild type and *cycd7;1-1* mutants. Graph shows the % of GMCs in S phase during a 90-minute incubation with EdU. Error bars indicate the 95% confidence interval. Asterisk indicates significant difference (p-value = 7x10E-6; Fisher’s Exact Test). Center lines in box plots show the medians; box limits indicate the 25th and 75th percentiles; whiskers extend 1.5 times the interquartile range from the 25th and 75th percentiles.

Next, we asked if mutations of *CYCD7;1* result in abnormal phenotypes. We obtained multiple alleles of *CYCD7;1:* FLAG_369E02 (*cycd7;1-1* (Collins et al., 2012), FLAG_498H08 (*cycd7;1-2*), GK_496G06- 019628, SALK_068526 and SALK_068526 (Fig. S2A). We determined by qRT-PCR that *cycd7;1-1* (FLAG_369E02) produced no transcript (Fig. S2B). On a whole plant level, we could not detect any abnormalities in *cycd7;1-1* compared to wild type (Fig. S1C). Because CYCDs promote G1/S transitions and CYCD7;1 is specifically expressed during the GMC divisions, we asked whether *cycd7;1-1* mutants halt this transition by counting GCs in cotyledons. Mutants in *cycd7;1-1* do not display fewer GCs compared to wild type 7 days after germination (dag) (Fig. S2D-F). However, at 4 dag, when cells in the earlier stages of the stomatal lineage are abundant, *cycd7;1-1* cotyledons have more GMCs compared to wild type cotyledons (Fig. 2D). Interestingly, the average size of *cycd7;1-1* GMCs is larger than wild type (Fig. 2E). We confirmed that these GMC abundance and size phenotypes were present in plants bearing a different allele of *CYCD7;1 (cycd7;1-2)* (Fig. S2G, H). Plant cells are known to increase in size during G1, so this phenotype suggests that CYCD7;1 hastens cell cycle progression in the GMC to GC transition. Because *cycd7;1-1* is the null allele, we characterized its phenotypes in more detail. We introgressed *pCDKB1;1:GUS,* which labels the transition from GMC to GCs (Boudolf et al., 2004), into *cycd7;1-1* mutants. Compared to wild type, *cycd7;1-1* mutants show increased number of GUS-positive cells suggesting that these cells remain longer in GMC fate before they divide into GCs (Fig. 2F-H). To directly test this hypothesis, we labeled S phases with 5-ethynyl-2’deoxyuridine (EdU) a thymidine analogue readily incorporated during DNA replication (Fig. 2I, J). Strikingly, significantly fewer GMCs in *cycd7;1-1* showed EdU labeling (indicating that they were in S phase during the EdU pulse) compared to wild-type GMCs (Fig. 2K). Together these data suggest that CYCD7;1 is required for GMCs to make a timely entry into S phase before their transition into GCs.

### CYCD7;1 interacts with RBR1

Typically, CYCDs drive the G1/S transition through inactivation of RBR1, and RBR1 activity was previously shown to be essential for repressing divisions in the stomatal lineage (Borghi et al., 2010; Matos et al., 2014). If CYCD7;1 and RBR1 function together, we would expect them to be co-expressed, to physically interact, and for there to be a phenotypic consequence of disrupting the interaction. Indeed, CYCD7;1 and RBR1 were shown to physically interact in BIFC and Y2H assays, dependent on the presence of the RBR1 binding motif L×C×E in CYCD7;1 (Matos et al., 2014). In addition, CYCD7;1 and RBR1 are co-expressed in GMCs (Fig. 3A-C). To test whether this interaction is functionally important, we took advantage of the fact that our translational reporter of CYCD7;1 triggers extra cell divisions in GCs (Fig. 1C, Fig. 3 D,E). Approximately 24% of GCs have one and 18% have two ectopic divisions in *pCYCD7;1:CYCD7;1-YFP* plants at 5 dag (Fig. 3G). If the RBR1 interaction is important for CYCD7;1 function, then mutation of the RBR1 binding motif LxCxE into LxGxK in CYCD7;1, should abrogate this division promoting activity. Strikingly, we found that *pCYCD7;1:CYCD7;1*^*LGK*^*-YFP* no longer triggers ectopic cell divisions in GCs (Fig. 3F,G). This effect was not due to differences in expression levels between *CYCD7;1-YFP* and *CYCD7;1*^*LGK*^*-YFP* (Fig S1B). Production of ectopic cell divisions in GCs, therefore, depends on the RBR1 binding residues in CYCD7;1.

**Figure 3:**
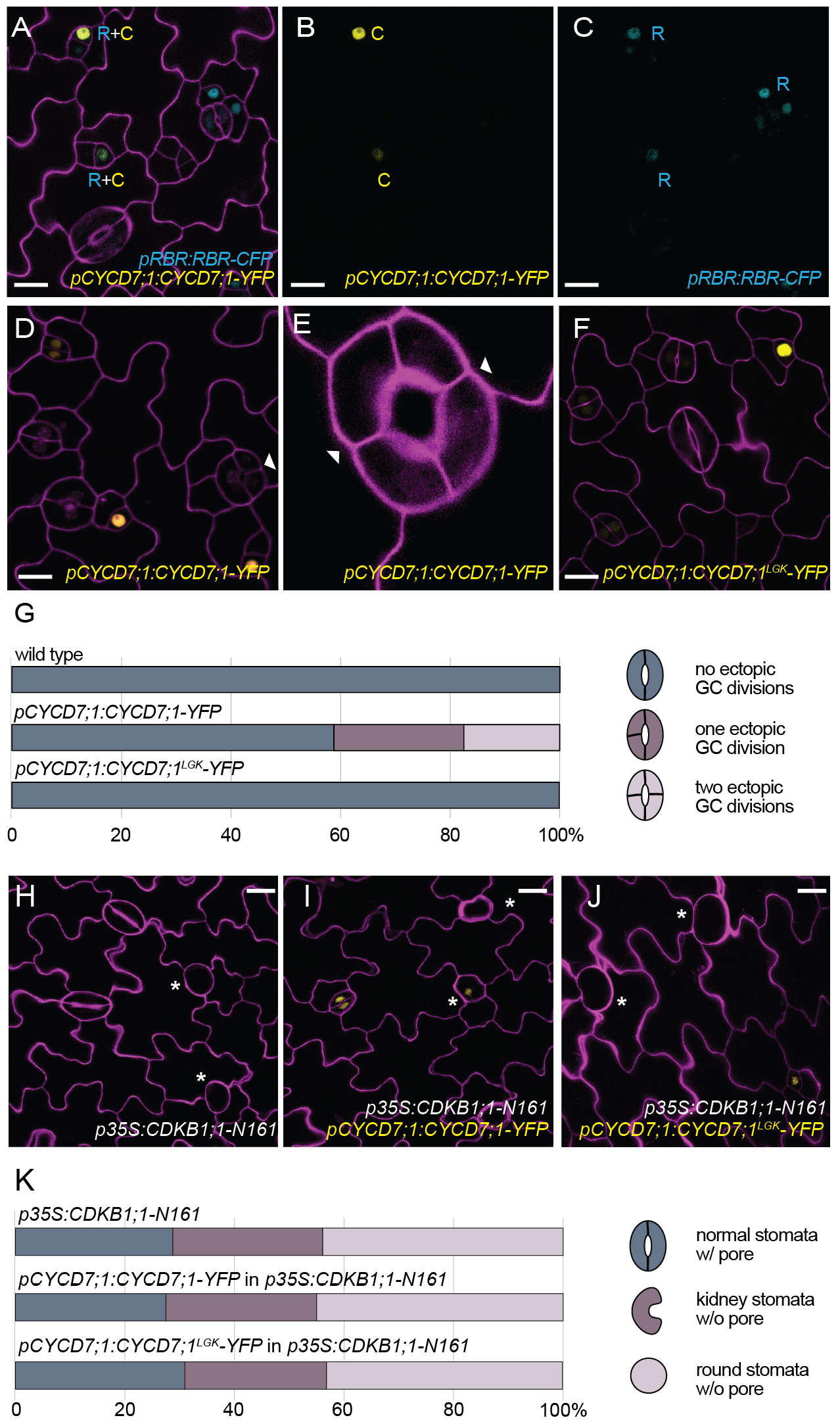
CYCD7;1 requires RBR1 binding and CDKB1;1 activity for ectopic cell divisions. **(A-C)** Co-expression of *pCYCD7;1:CYCD7;1-YFP* (yellow, C) and *pRBR1:RBR1-CFP* (cyan, R) in GMCs at 5 dag. **(D-E)** Expression of *pCYCD7;1:CYCD7;1-YFP* drives ectopic cell divisions (white arrowheads). **(F)** Expression of *pCYCD7;1:CYCD7;1^LGK^-YFP* (yellow) does not drive ectopic cell divisions. **(G)** Quantification of ectopic cell divisions in GCs at 5 dag in cotyledons in wild type (N=173), *pCYCD7;1:CYCD7;1-YFP* (N=306) and *pCYCD7;1:CYCD7;1^LGK^-YFP* (N=288). **(H)** Phenotype of dominant negative *p35S:CDKB1;1-N161* at 6 dag. White asterisks label arrested GMCs. **(I-J)** Failure of *pCYCD7;1:CYCD7;1-YFP* **(I)** and *pCYCD7;1:CYCD7;1*^*LGK*^*-YFP* (J) to suppress *CDKB1;1-N161* phenotype at 6 dag. White asterisks label arrested GMCs. **(K)** Quantification of stomata phenotypes in cotyledons in *p35S:CDKB1;1-N161* (N=238), *pCYCD7;1:CYCD7;1-YFP* in *o35S:CDKB1;1-N161* (N=296) and *pCYCD7;1:CYCD7;1*^*LGK*^*-YFP* in *p35S:CDKB1;1-N161* (N=217) at 6 dag. Confocal images show cell outlines (magenta) stained with propidium iodide. Scale bar 10 μm (A-D, F) and 20 μm (H-J).

### CYCD7;1 needs CDKB1 activity to drive ectopic divisions

Cyclins bind to CDKs to ensure kinase activity and completion of cell division; undivided cells expressing GC fate markers result from reduction or loss of CDK activity (e.g., hypomorphic *cdka;1* mutants (Weimer et al., 2012), *cdkb1;1 cdkb1;2* double mutants (Xie et al., 2010) or dominant-negative CDKB1;1-N161 (Boudolf et al., 2004)). To test whether CYCD7;1 required CDK activity to drive divisions, we expressed CYCD7;1-YFP and CYCD7;1^LGK^-YFP under the CYCD7;1 promoter in plants bearing a dominant negative version of *CDKB1;1* (CDKB1;1-N161, Fig. 3H-J). Although we could see expression of both CYCD7;1 markers in arrested GMCs, they could neither rescue the phenotype nor trigger ectopic cell divisions (Fig. 3I-K). Thus CYCD7;1 requires CDKB1 activity either as a partner, or downstream at the G2/M transition for completion of the division.

### CYCD7;1 expression domain is constrained by stomatal lineage transcription factors

Our evidence points to CYCD7;1 acting like a canonical CYCD, therefore we turned our attention to regulation of its highly restricted expression pattern. Three transcription factors are contemporaneously expressed with CYCD7; 1—MUTE, FAMA and FLP (Fig 1I-K)—but MUTE precedes CYCD7;1 while the others persist longer. Given these patterns, we tested whether MUTE was necessary for CYCD7;1 expression. When *pCYCD7;1:CYCD7;1-YFP* was crossed into the *mute* mutant, we could observe the typical *mute* phenotype of many small meristemoid-like cells that fail to differentiate into GMCs (Pillitteri et al., 2007). In a few of these meristemoid-like cells, we detected weak CYCD7;1-YFP signal (Fig. 4A,B). Fluorescence intensity measurements showed that CYCD7;1-YFP signals in *mute* are ~50% reduced (Fig 4C-F) indicating that MUTE promotes CYCD7;1 expression, though it is not absolutely essential for it. In none of these images did we observe any ectopic divisions of the meristemoid-like cells.

**Figure 4:**
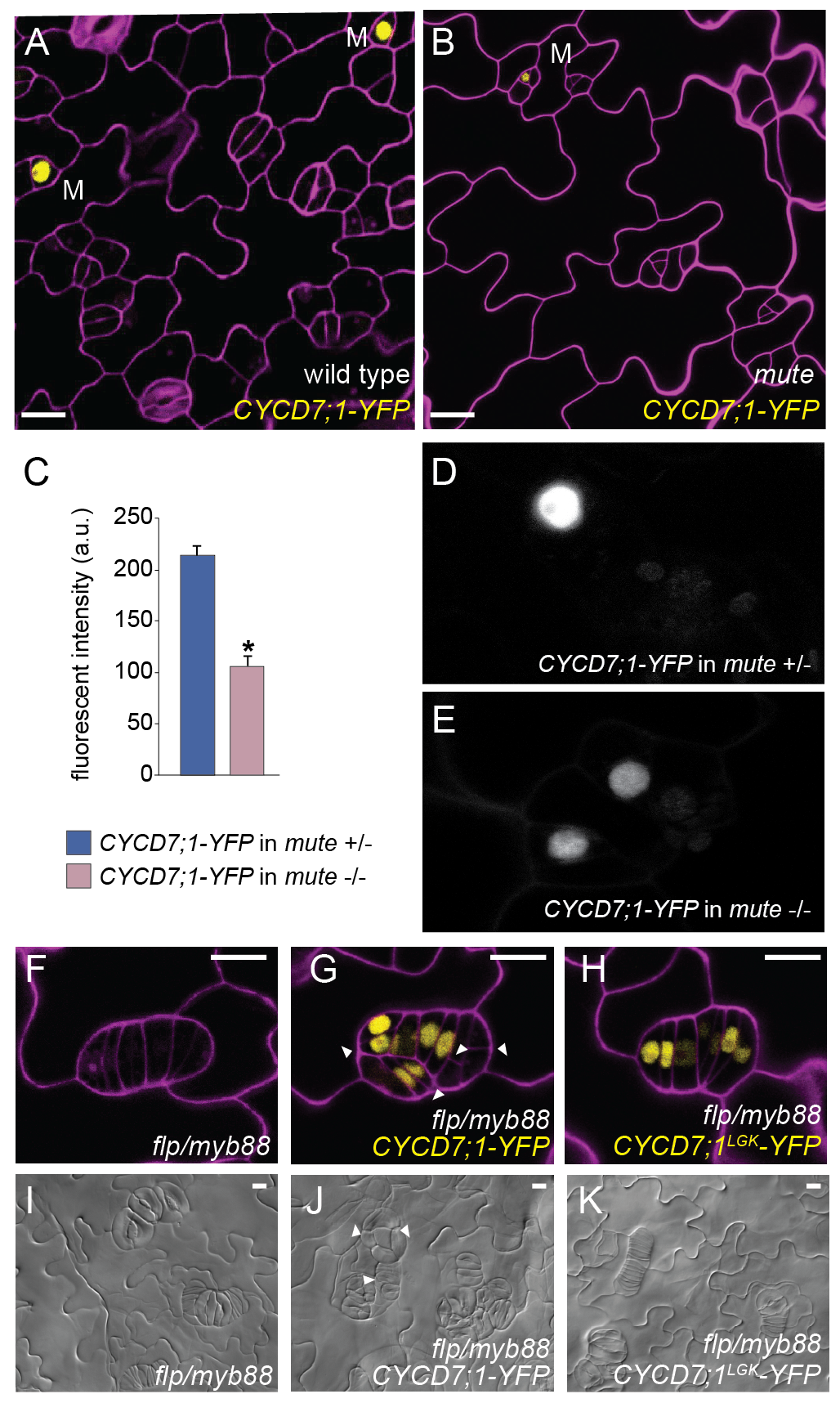
CYCD7;1-YFP is expressed at low levels in *mute* mutants and persists and drives ectopic divisions in *flp/myb88* mutants. **(A, B)** Wild type and *mute* mutants expressing *pCYCD7;1:CYCD7;1-YFP* in 6 day old cotyledons. Scale bar 10 μm; M, meristemoid. (C-E) Quantification of fluorescence intensity of CYCD7;1-YFP in homozygous *mute* mutants (N=27) and their heterozygous or wild-type sister plants (N=21) (a.u., arbitrary units). Images of cotyledons were taken at 4 dag. Error bars show standard error. Asterisk shows statistical significance (p-value <0.0001; Student-t test). **(F)** Phenotype of the double mutant *flp/myb88* at 6 dag. **(G)** Expression of *pCYCD7;1:CYCD7;1-YFP* in *flp/myb88* drives ectopic divisions in tumors at 6 dag. (H) Expression of *pCYCD7;1:CYCD7;1*^*LGK*^*-YFP* in *flp/myb88* is less able to drive ectopic divisions at 6 dag. **(I)** DIC images of the phenotype of the double mutant *flp/myb88* at 12 dag. **(J)** Expression of *pCYCD7;1:CYCD7;1-YFP* in *flp/myb88* drives ectopic divisions in tumors at 12 dag. **(K)** Expression of *pCYCD7;1:CYCD7;1*^*LGK*^*-YFP* in *flp/myb88* is less able to drive ectopic divisions at 12 dag. White arrowheads label ectopic divisions. Confocal images show cell outlines (magenta) stained with propidium iodide. Scale bar 10μM.

CYCD7;1 appears to be repressed during FAMA’s expression peak. We therefore tested whether FAMA, in its role as the master transcriptional regulator of stomatal division and differentiation, is a direct regulator of CYCD7;1. In *fama* mutants GMCs divide repeatedly without attaining GC fate (Fig. 5A-E) and these “tumors” express CYCD7;1-YFP (Fig. 5B,C); although the reporter fades in older leaves suggesting that CYCD7;1-YFP is also subject to posttranslational regulation (Fig. 5D,E). In the *fama* tumors, *pCYCD7;1:CYCD7;1-YFP* drives ectopic divisions (Fig. 5B,D, white arrowheads), but the *CYCD7;1*^*LGK*^ version that cannot bind RBR1, does not (Fig. 5C,E). To test whether FAMA might directly regulate CYCD7;1, we extracted reads from a FAMA ChIP-seq experiment, performed under similar conditions as in (Lau and Bergmann, 2015; Lau et al., 2014). As shown in Fig. 5F, it is clear that FAMA is associated with the promoter region and gene body of *CYCD7;1*.

**Figure 5:**
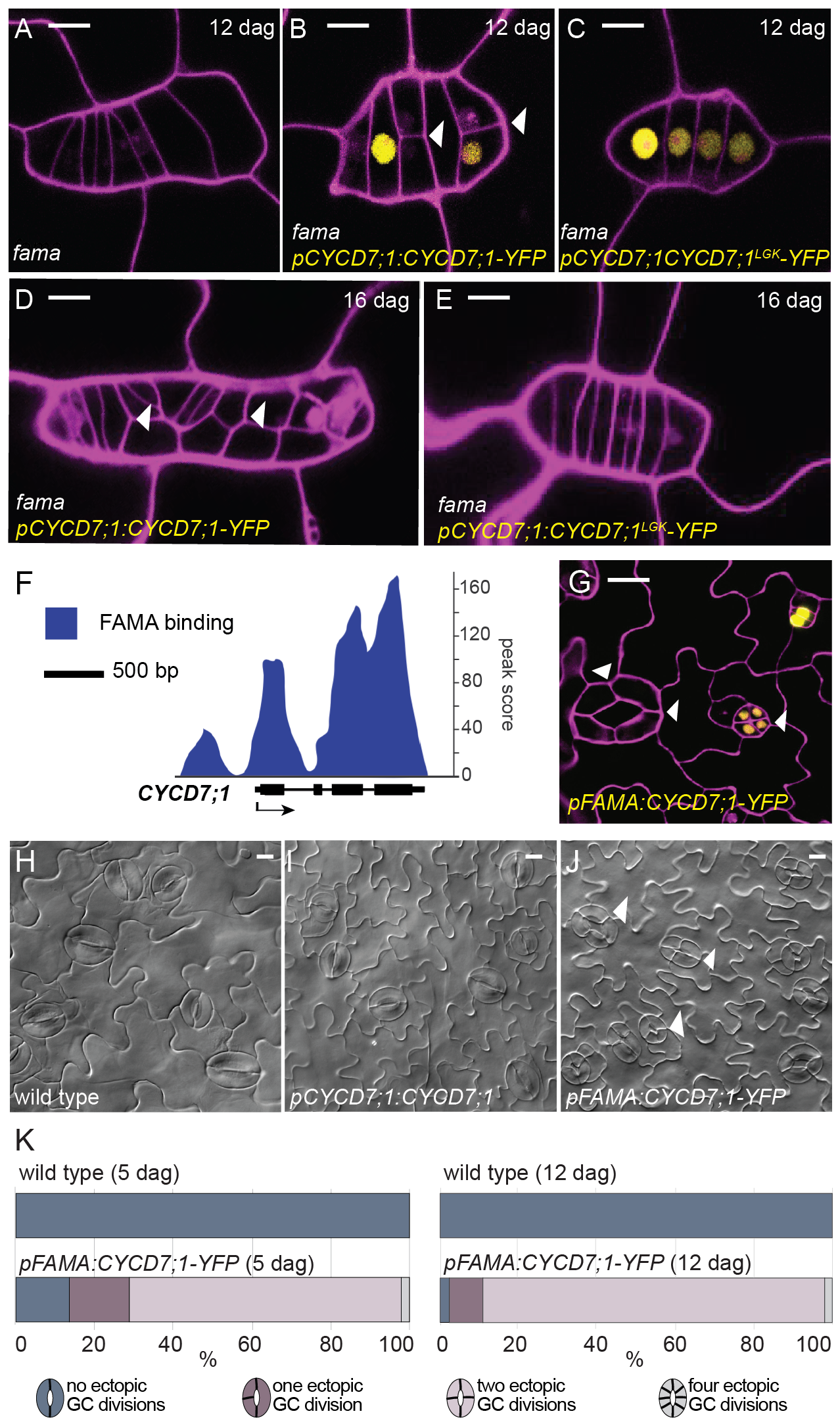
CYCD7;1 expression is regulated by FAMA which serves to constrain CYCD7;1 activity. **(A-E)** Confocal images of *fama, pCYCD7;1:CYCD7;1-YFP* in *fama* mutant background and *pCYCD7;1:CYCD7;1*^*LGK*^*-YFP* in *fama* mutant background at 12 or 16 dag, respectively. **(F)** ChlP-Seq profile of FAMA binding to the promoter and gene body of *CYCD7;1.* Black arrow indicates gene orientation and transcriptional start sites. **(G)** Confocal image of *pFAMA:CYCD7;1-YFP* at 5 dag. White arrowheads show ectopic division and prolonged CYCD7;1-YFP presence. **(H-J)** DIC images of abaxial cotyledon epidermis of wild type, *pCYCD7;1:CYCD7;1* and *pFAMA:CYCD7;1-YFP* at 12 dag. Scale bar, 10μM. Arrowheads point at ectopic cell divisions. (K) Quantification of ectopic cell divisions in wild type (N=142) and *pFAMA:CYCD7;1-YFP* (N=237) at 5 dag and in wild type (N=125) and *pFAMA:CYCD7;1-YFP* (N=153) at 12 dag. Confocal images show cell outlines (magenta) stained with propidium iodide. Scale bar 10μm.

Along with FAMA, two partially redundant R2R3 MYB transcription factors, FOUR LIPS (FLP) and MYB88, restrict GMC divisions. Previously, it was shown that FLP/MYB88 bind directly to the *CDKB1;1* promoter and can repress *CDKB1;1* transcription (Lee et al., 2013; Vanneste et al., 2011; Xie et al., 2010). *flp/myb88* mutants also display GMC overproliferation but, unlike *fama* mutants, some differentiated GCs form (Lai et al., 2005; Xie et al., 2010), Fig. 4F,I). *CYCD7;1-YFP* (and *CYCD7;1*^*LGK*^*-YFP*) translational reporters are highly expressed in *flp/myb88*, and CYCD7;1-YFP, but not CYCD7;1^LGK^_YFP, induces ectopic divisions (Fig. 4 G,H,J,K).

The phenotypes of loss and gain of CYCD7;1 activity suggest that its narrow window of expression is essential to guarantee a 2-celled stomatal complex. Using the *FAMA* promoter in wild type, thus driving CYCD7;1 slightly later than under its endogenous cis-regulatory control, we find a dramatic enhancement of ectopic divisions (Fig. 5G-K). Compared to *pCYCD7;1:CYCD7;1-YFP* in which ~24% of stomata were four-celled at 5 dag, in *pFAMA:CYCD7;1-YFP*, that number was ~70%, with 2% of stomata being 8-celled (N=237). The amount of four-celled stomata increases to 87% at 12 dag, with another 2% being 8-celled (N=153). (Fig. 5K). Quantification of fluorescence intensity indicates that expression with *FAMA* and *CYCD7* promoters yields equivalent levels of CYCD7;1-YFP in GMCs (Fig S1B), however, this fusion protein persists in ectopically divided GCs when expressed under the FAMA promoter (Fig. 5L). This directly links the activity of FAMA as a lineage specific transcription factor with the cell cycle regulator CYCD7;1 to ensure “one and only one division” to create a pair of guard cells.

## Discussion

We have shown that CYCD7;1 is specifically expressed in GMCs prior to the last symmetric cell division that forms the 2-celled stomatal complex. Depletion of *CYCD7;1* slows down this cell division whereas ectopic expression of CYCD7;1 can trigger cell divisions in GCs. Mutation of the RBR1 binding motif in CYCD7;1 disrupts its interaction with RBR1 and renders CYCD7;1^LGK^ incapable of driving ectopic division. The connection to RBR1 fits with previous work showing that CYCD7;1 interacts with CDKA;1 (Van Leene et al., 2010), together supporting a role for CYCD7;1 in the canonical regulatory complex for G1/S transitions and the commitment to divide. CYCD7;1 activity in cell cycles, however, is directly repressed by the lineage specific transcription factor FAMA to ensure a coupling between the cell division which terminates the stomatal lineage, and the formation of terminally fated GCs. This interconnection represents a direct link between cell cycle regulators and developmental decisions (Fig. 6).

**Figure 6:**
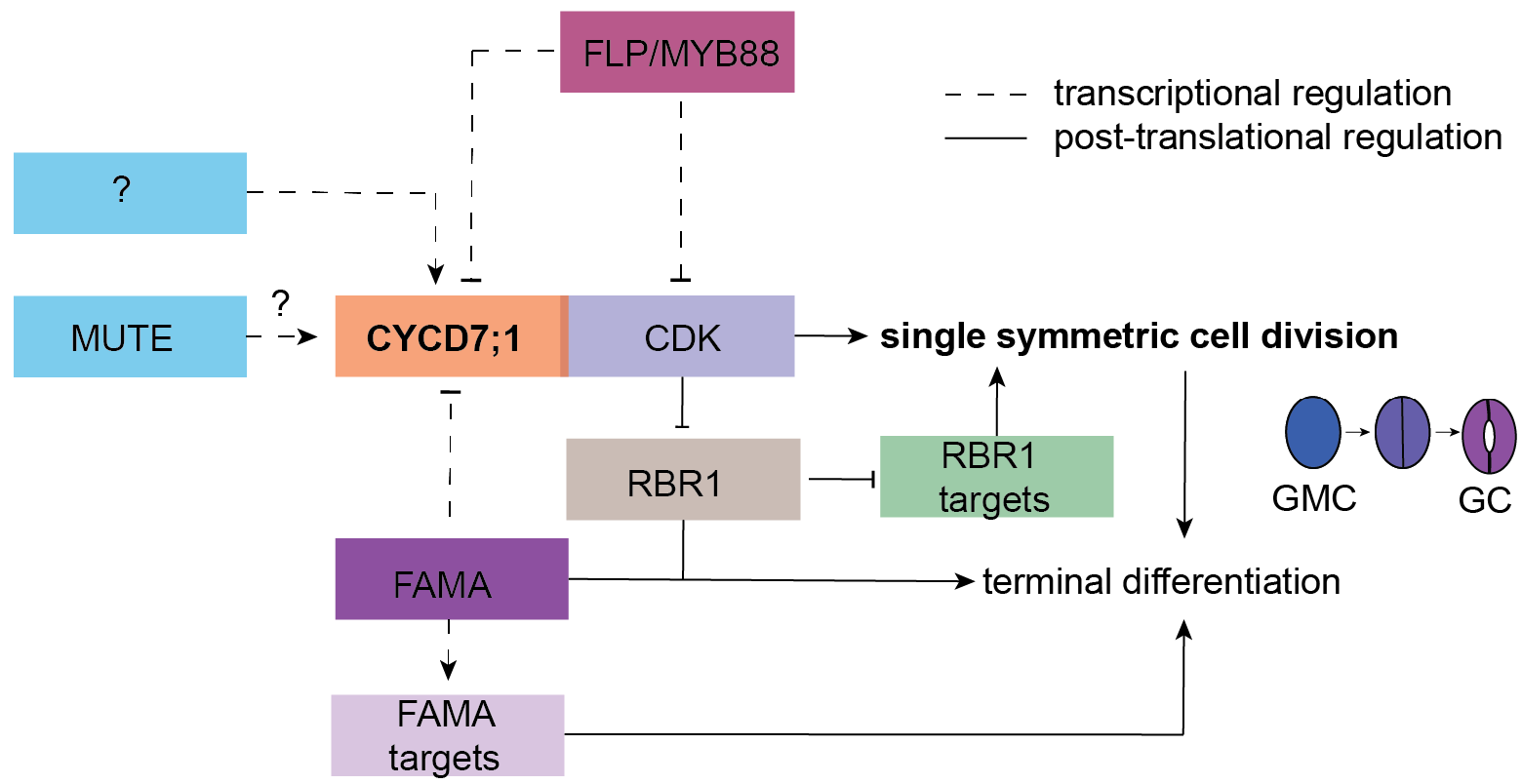
Model of the developmental integration of CYCD7;1 to ensure lineage specific cell cycle regulation. Cell cycle regulators are integrated with stomatal specific transcriptions factors to ensure the last formative division of the lineage that creates one pair of symmetric guard cells. Initiation of CYCD7;1’s expression in GMCs requires factors in addition to MUTE (question mark). CYCD7;1 together with its CDK partner executes the formative division of the GMC. Due to the observation that this last division is not completely abolished in *cycd7;1* mutants, other D-type cyclins likely back up G1-S phase transition. CDK/CYCD complexes phosphorylate RBR1 in order to release its negative function on S phase promoting factors. To ensure termination of the lineage, the transcription factor FAMA, itself slightly later expressed than CYCD7;1, binds to the CYCD7;1 promoter to temporally control expression of the lineage-specific CYCD7;1 to GMCs and to restrict the cell cycle right after the last division. Transcriptional regulation is marked by dashed lines. This regulatory network ensures high cell cycle activity for the last formative division in the stomatal lineage and terminates cell division activity to “one and only one” division.

CYCDs are critical for the G1/S transition and commitment to divide, and are therefore interesting candidate hubs for the integration of developmental control with the cell cycle machinery. In *Arabidopsis,* there are 10 D-type cyclins, some active in multiple tissues (CYCD3s, CYCD4s, CYCD2;1) but others whose activity is linked to specific cell types (CYCD6;1 and CYCD7;1) or cell cycle behaviors (CYCD5;1 endoreplication) (Dewitte et al., 2007; Kono et al., 2007; Sanz et al., 2011; Sterken et al., 2012) (Adrian et al., 2015; Sozzani et al., 2010), this study). Phylogenetic analyses showed that CYCD6;1 and CYCD7;1 proteins diverge from other D-type cyclins in Arabidopsis (Wang et al., 2004), but also that CYCD7;1 most closely resembles the single D-type cyclin in *Physcomitrella* (Menges et al., 2007), consistent with our observation that it could promoting G1/S transitions (a core D-type activity) in multiple cell types.

Interestingly, both CYCD6;1 and CYCD7;1 are limiting for essential formative divisions during development. In the root, CYCD6;1 is important for the cortex endodermis initial daughter (CEID) cell divisions (Sozzani et al., 2010; Weimer et al., 2012). Here, SHORTROOT (SHR) directly activates expression of CYCD6;1 which works in concert with CDKA;1 to trigger the formative division of the CEID (Cruz-Ramírez et al., 2012; Sozzani et al., 2010; Weimer et al., 2012). This interaction promotes the initiation of an asymmetric stem-cell division program. In contrast, CYCD7;1 expression marks the boundary between two types of divisions: the continual asymmetric divisions of meristemoids vs. the single symmetric division of a GMC. Here we find a quantitative requirement for *MUTE* to promote full CYCD7;1 expression, but a clear requirement for FAMA and FLP/MYB88 to repress CYCD7;1 after GMC division. The low expression level of CYCD7;1 in the absence of *MUTE* may point to a direct role for *MUTE* in activating CYCD7;1 expression. MUTE is structurally similar to FAMA, and therefore might be able to interact with *CYCD7;1* regulatory sequences. Alternatively, as meristemoid cells in *mute* never transition into GMCs, low *CYCD7;1* levels may be an indirect consequence of altered cell fate. In either case, it is notable that the introduction of CYCD7;1-YFP in *mute* did drive not additional meristemoid cell divisions suggesting that CYCD7;’s division-promoting behavior requires a threshold level not reached in this genetic background.

It is tempting to speculate that spatiotemporal restriction of CYCDs could be a mechanism to control the cell cycle machinery more efficiently and to cope with different developmental programs. The importance of these specialized CYCDs, however, must be squared with the relatively minor phenotypes associated with their loss—neither *CYCD7;1* nor *CYCD6;1* mutants abolish the production of specialized cells or tissue layers (Fig. 2) (Sozzani et al., 2010)). Most likely, CYCD6;1 and CYCD7;1 assist other, more general, cyclins in executing the cell division programs or ensure particularly high cell cycle kinase activity. In the case of the stomatal lineage, CYCD3;1 and CYCD3;2, despite being considered general G1/S cyclins (Dewitte et al., 2007; Dewitte et al., 2003; Menges et al., 2006), also show high expression in the stomatal lineage (Adrian et al., 2015). It is also important to recognize that CYCD/CDKA complexes likely have many downstream targets and that increased kinase activity could induce different downstream processes, either in a feedback loop or for differentiation processes. In plants, specific CDK/cyclin complexes can have differential activity towards individual substrates, and both CDK and cyclin proteins contribute to substrate recognition (Harashima and Schnittger, 2012), however, there is evidence that between the CDK and cyclin, the cyclin may have a more prominent role (Weimer et al., 2016). Specific expression of individual cyclins, such as CYCD7;1 in the stomatal lineage, therefore, could contribute to fine-tuning of cell division control and downstream substrate recognition.

Leaves lose overall division competency as they mature, leading to a situation where GMCs are surrounded by post-mitotic cells. Formation of functional stomata, however, requires a cell division to produce two cells, suggesting that this division has unique additional regulation. Stomata are found in remarkably diverse patterns and exhibit a 10-fold variation in size in different species (McElwain et al., 2016), yet there have still to be reports of more than two stomatal guard cells flanking a pore. Therefore, despite the ease with which we could create four-celled stomata through experimental manipulation, in nature, regulation to ensure a single division appears crucial.

## Material and Methods

### Plant material and growth conditions

*Arabidopsis thaliana* Columbia-0 (Col-0) was used as wild type in all experiments. All mutants and transgenic lines tested have this ecotype background. Seedlings were grown on half-strength Murashige and Skoog (MS) medium (Caisson labs, USA) medium at 22°C under 16 hour-light/8 hour-dark cycles and were examined at the indicated time. The following previously described mutants and reporter lines were used in this study: *mute* (Pillitteri et al., 2007); *fama-1* (Ohashi-Ito and Bergmann, 2006); *flp;myb88* (Lai et al., 2005); *proSPCH:SPCH:CFP* and *proMUTE:MUTE-YFP* (Davies and Bergmann, 2014); *proRBR1:RBR1-CFP* (Cruz-Ramírez et al., 2012), *pro35S:CDKB1;1-N161* (Boudolf et al., 2004); *proCDKB1;1:GUS* (Boudolf et al., 2004).

### *CYCD7;1* mutants

CYCD7;1 mutants FLAG_369E02 *(cycd7;1-1)* and FLAG_498H08 *(cycd7;1-2)* were derived from the INRA/Versaille collection (Versaille, France) and *cycd7;1;1* was backcrossed twice to Col-0. GK_496G06-019628 was derived from the GABI-Kat collection (Cologne, Germany). SALK_068423 and SALK_068526 were obtained from ABRC (Columbus, USA).

### Vector construction and plant transformation

Constructs were generated using the Gateway® system (Invitrogen, CA, USA). Appropriate genome sequences (PCR amplified from Col-0 or from entry clones) were cloned into Gateway compatible entry vectors, typically pENTR/D-TOPO (Life Technologies, CA, USA), to facilitate subsequent cloning into plant binary vectors pHGY (Kubo et al., 2005) or R4pGWB destination vector system (Nakagawa et al., 2008; Tanaka et al., 2011). The translational reporter for CYCD7;1 was generated by cloning the genomic fragment (promoter+CDS) into the entry vector pENTR to generate the entry vector CYCD7;1-genomic-pENTR, followed by LR recombination into the destination vector pHGY to generate the final construct. For the translational reporter for CYCD7;1^LGK^, the LxCxE motif of CYCD7;1-genomic-pENTR was mutated to L×G×K by site directed mutagenesis using the QuikChange II Kit (Agilent, CA, USA) to generate the entry clone CYCD7;1-genomic-pENTR and then recombined into pHGY. The transcriptional reporters for CYCD7;1 were generated by cloning the CYCD7;1 promoter region into pENTR, then recombined into the destination vectors pHGY (cytosolic YFP). The other constructs generated in this study *proCYCD7;1:YFP-YFPnls, proFAMA:FAMA-CFP, proML1:CYCD7;1-YFP, proML1:CYCD7;1, proCYCD7;1:CYCD7;1*, and *proFAMA:CYCD7;1-YFP* were generated with the tripartite recombination of the plant binary vector series R4pGWB (Nakagawa et al., 2008; Tanaka et al., 2011), with the Gateway entry clones of the promoters and coding sequences compatible with the binary R4pGWB destination vector system. Primer sequences used for entry clones are provided in Table 1. Transgenic plants were generated by Agrobacterium–mediated transformation (Clough, 2005) and transgenic seedlings were selected by growth on half-strength MS plates supplemented with 50 mg/L Hygromycin (pHGY, p35HGY, pGWB1, pGWB540 based constructs) or Kanamycin 100 mg/L (pGWB440 and pGWV401 based constructs) or 12 mg/L of Basta (pGWB640 based constructs).

### Confocal and DIC microscopy

For confocal microscopy, images were taken with a Leica SP5 microscope and processed in ImageJ. Cell outlines were visualized by either 0.1 mg/ml propidium iodide in water (Molecular Probes, OR, USA) incubation for 10 min, rinsed in H_2_O once). For DIC microscopy, samples were cleared in 7:1 ethanol:acetic acid, treated 30 min with 1N potassium hydroxide, rinsed in water, and mounted in Hoyer's medium. Differential contrast interference (DIC) images were obtained from the middle region of adaxial epidermis of cotyledons on a Leica DM2500 microscope or Leica DM6 B microscope.

### Quantification of fluorescent intensity

Images of GMCs in cotyledons were taken at 4 dag with identical settings and processed in ImageJ. Fluorescent intensity was measured as mean gray value in the nucleus, subtracted by the background. Measurements were averaged for mutant and control experiments with Student’s-t-test used to determine the statistical significance.

### GUS staining

5-day old seedlings were incubated in staining solution for 12 hours and destained in 70% ethanol at 60–70°C for four hours. Staining solution for 5ml: 100μl of 10% Triton X-100, 250μl 1M NaPO4 (pH 7.2), 100μl 100mM potassium ferrocyanide, 100μl potassium ferricyanide, 400μl 25 mM X-Gluc, 4050μl dH2O. Images were taken with a Leica DM6 B microscope.

### EdU labeling

EdU labeling was performed using the Click-iT® EdU Alexa Fluor® 488 Imaging Kit (ThermoFisher Scientific, MA, USA). 4-day old seedlings were incubated in 20μM EdU solution in half-strength MS for 90 minutes at room temperature. Seedlings were transferred to new tubes and washed three times with wash buffer (1% BSA in PBS). Wash buffer was removed and fixation buffer was added (3.7% formaldehyde in PBS) for 30 min at room temperature. Seedlings were transferred to new tubes and washed two times with permeabilization buffer (0.5% Triton x-100 in PBS) for 10 minutes each, protected from light on a slow rocking platform. Plants were transferred to new tubes and incubated in reaction cocktail (455μL Click-IT reaction buffer, 20μL CuSO_4_, 2μL Alexa Fluor Azide 488, 25 μL 1x Click-IT EdU additive) for 1 hour at room temperature, protected from light, without agitation. Seedlings were transferred to new tubes and washed twice for 10 minutes at room temperature with wash buffer on a slow rocking platforms, protected from light. Cotyledons were imaged using a Leica SP5 microscope not more than two hours after the completion of washes and processed in ImageJ.

### qPCR

100 mg ground frozen material from 8-day old plants was used for RNA extraction according to the manufacture’s manual (RNeasy Mini Kit, Qiagen, Germany). 1μg total RNA was used as a template for cDNA synthesis (iScript cDNA synthesis kit, BioRad, CA, USA). qPCR setup was according to the manual of the SsoAdvanced Universal SYBR Green Supermix (BioRad, CA, USA). qPCR was performed by CFX96 Real Time C1000 Thermal Cycler (BioRad, CA, USA) according to the following reaction conditions: 95°C for 30 s, followed by 39 cycles at 95°C for 10 s and at 60°C for 30 s. ACTIN was used as a reference gene for all qPCRs performed. Primers can be found in Table 1.

**Table 1:**
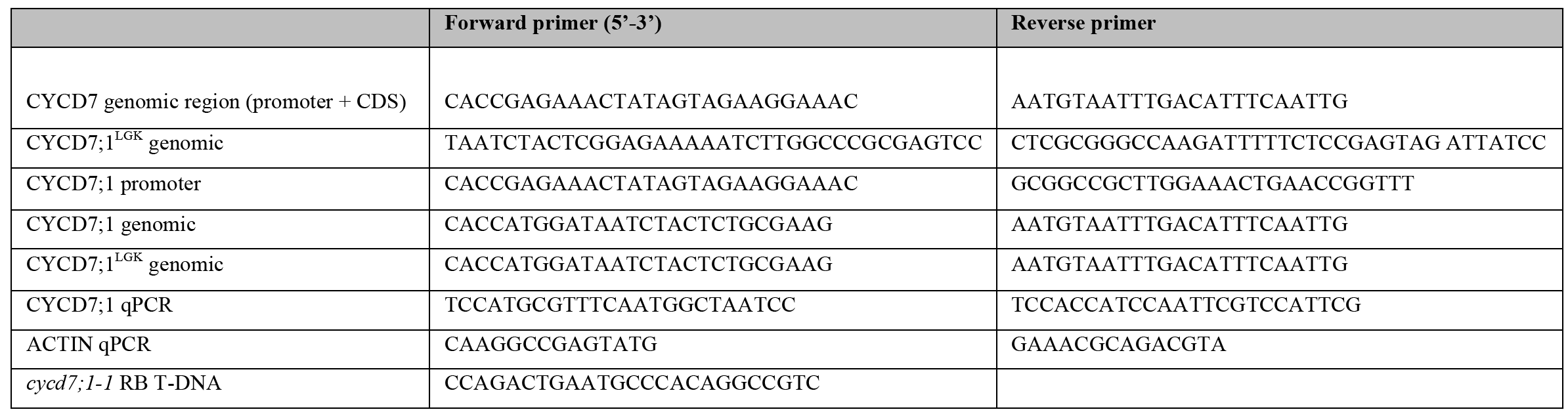
Primers used in this study.

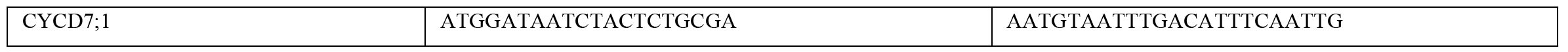

## Acknowledgments

We thank members of the Bergmann lab for helpful comments on the manuscript and Charles Hachez (UCLouvain) for his contributions to the initiation of the CYCD7;1 project.

## Supplementary Figures

**Figure S1:**
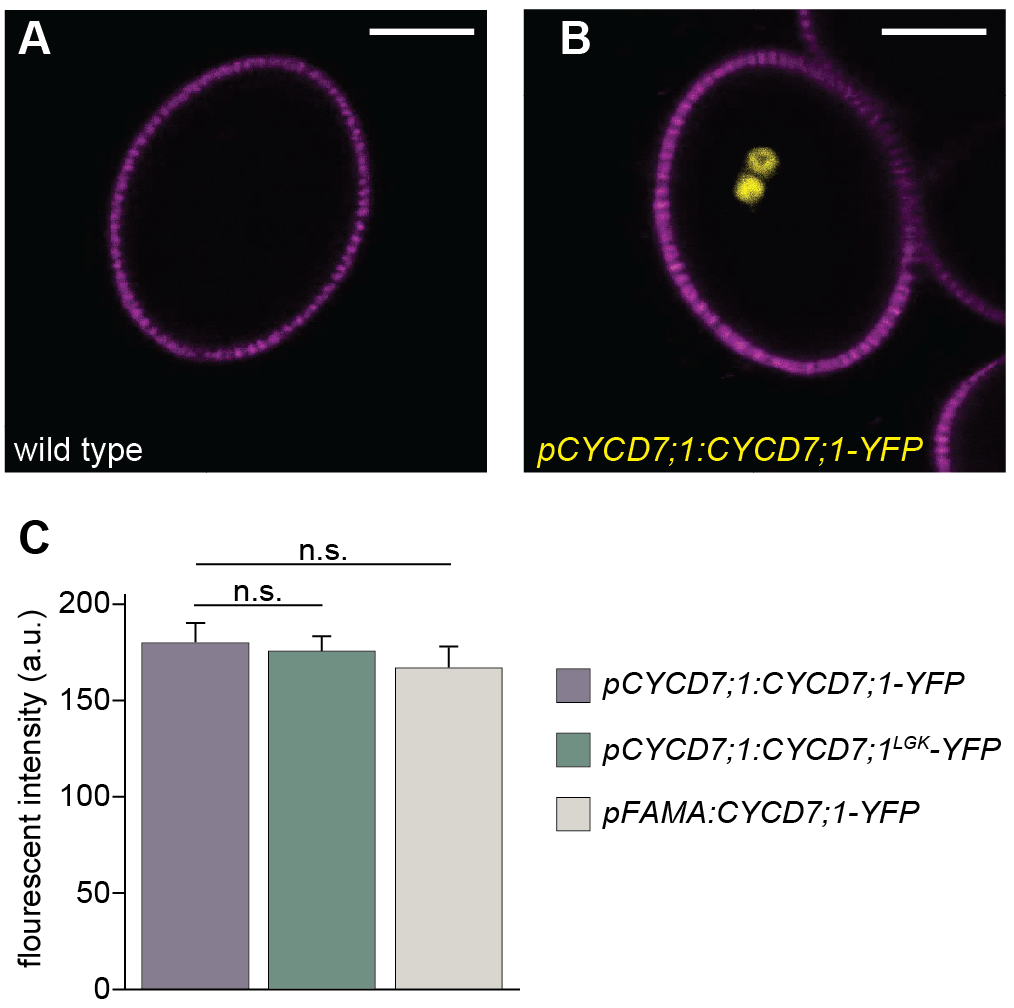
CYCD7;1 expression patterns. **(A, B)** CYCD7;1 (yellow) is expressed in sperm cells during pollen anthesis. (C) Intensity measurements of fluorescent nuclei were 179 a.u. +/-10 SE for *proCYCD7;1:CYCD7;1-YFP* vs 176 a.u. +/-8 SE for *proCYCD7;1:CYCD7;1*^*LGK*^*-YFP* (N=15 nuclei/line; p> 0.05; Student’s t-test) and *proCYCD7;1:CYCD7;1-YFP* 166 a.u. +/−11 SE for *proFAMA:CYCD7;1-YFP* (N=15 nuclei/line; p> 0.05; Student’s t-test). Error bars show standard error. a.u., arbitrary units; n.s. non-significant; SE, standard error.

**Figure S2:**
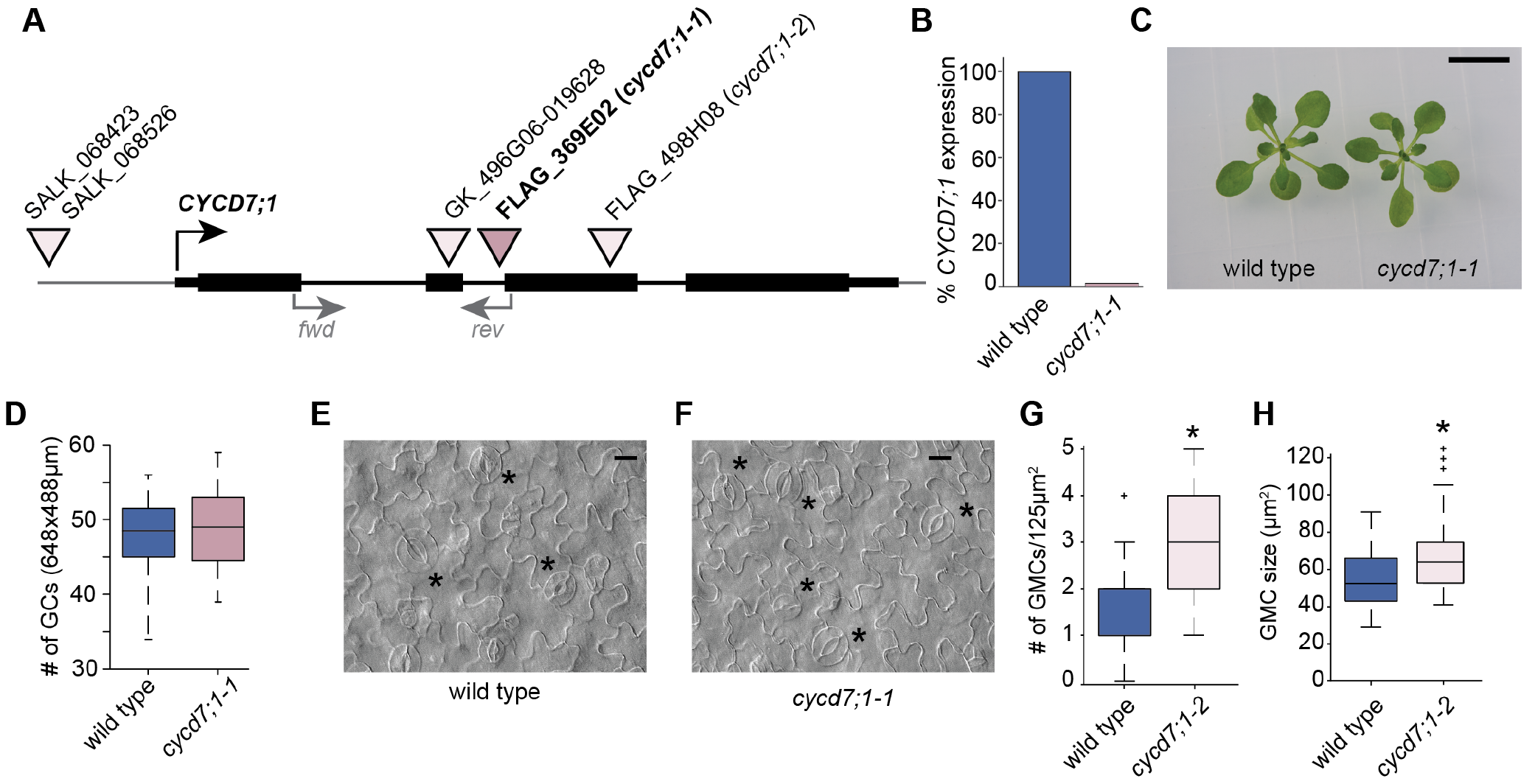
T-DNA insertion lines and phenotype of *cycd7;1* mutants. **(A)** Schematic drawing of *CYCD7;1* gene structure with available T-DNA insertion lines and their insertion sites. Black boxes indicate exons. Gray arrowheads marked with *fwd* and *rev* show primer binding sites for qPCR. **(B)** qPCR of *CYCD7;1* expression in wild type and the *cycd7;1-1* mutant. Primer binding sites are shown in (A). **(C)** Wild type and *cycd7;1-1* mutant seedlings at 14 dag. (D) Quantification of GCs in wild type and *cycd7;1-1* mutants at 5 dag on the abaxial side of cotyledons (N =12 cotyledons for each genotype). Difference between the wild type and *cycd7;1-1* is not significant (p-value = 0.8169; Mann-Whitney U test). **(E)** Wild type cotyledon with mature GCs, labeled with black asterisks at 7 dag. **(F)** Cotyledon of *cycd7;1-1* mutant with mature GCs, labeled with black asterisks, images were taken at 7 dag. **(G)** Quantification of the number of GMCs in wild type and *cycd7;1-2* cotyledons at 4 dag. Asterisk indicates significant difference (p-value = 0.0031; Mann-Whitney U test). **(H)** Quantification of GMC area in wild type (N=29) and *cycd7;1-2* (N=46) cotyledons, 4 dag. Asterisk indicates significant difference (p-value = 0.0053; Mann-Whitney U test). Center lines show the medians; box limits indicate the 25th and 75th percentiles; whiskers extend 2.5 times the interquartile range from the 97.5th percentile. Scale bar 1 cm in (C) and 20 μM in (E and F).

